# Recurrent losses and rapid evolution of the condensin II complex in insectsg

**DOI:** 10.1101/471243

**Authors:** T. King, C.J. Leonard, J.C. Cooper, S. Nguyen, E. Joyce, N. Phadnis

## Abstract

Condensins play a crucial role in the organization of genetic material by compacting and disentangling chromosomes. The condensin I and condensin II complexes are widely considered to have distinct functions based on studies in a few model organisms, although the specific functions of each complex are yet to be fully understood. The condensin II complex is critical for genome organization in Drosophila, and is a key anti-pairing factor that separates homologous chromosomes in somatic cells. Intriguingly, the Cap-G2 subunit of condensin II is absent in *Drosophila melanogaster,* and this loss may be related to the high levels of homologous chromosome pairing in somatic cells seen in flies. Here, we find that this Cap-G2 loss predates the origin of Dipterans, and other CapG2 losses have occurred independently in multiple insect lineages. Furthermore, the Cap-H2 and Cap-D3 subunits have also been repeatedly and independently lost in several insect orders, and some taxa lack condensin II-specific subunits entirely. We used Oligopaint DNA-FISH to quantify pairing levels in ten species across seven orders, representing several different configurations of the condensin II complex. We find that all non-Dipteran insects display near-uniform low pairing levels, suggesting that some key aspects of genome organization are robust to condensin II subunit losses. Finally, we observe consistent signatures of positive selection in condensin II subunits across flies and mammals. These findings suggest that these ancient complexes are far more evolutionarily labile than previously suspected, and are at the crossroads of several forms of genomic conflicts. Our results raise fundamental questions about the specific functions of the two condensin complexes and the interplay between them in taxa that have experienced subunit losses, and open the door to further investigations to elucidate the diversity of molecular mechanisms that underlie genome organization across various life forms.

## Introduction

All living organisms must organize their genetic material, and the molecular machinery involved in the fundamental processes of genome organization is deeply conserved. Condensins are key players in the tasks of compacting and disentangling chromosomes to ensure proper segregation of genetic material (1, 2). All cellular life on Earth, including bacteria, archea, fungi, plants and animals, possess condensins or an analogous protein complex (3, 4). The function of condensins is so integral to chromosome mechanics that they are thought to have arisen before histones (5).

Two distinct condensin complexes are present in most multicellular eukaryotes. Condensin I and condensin II are pentameric complexes that share a hinge structure made up of the SMC2 and SMC4 subunits. The non-SMC subunits of condensin I consist of the kleisin Cap-H, which serves a scaffold and linker, and is bound by the HEAT-repeat subunits Cap-D2 and Cap-G. Analogously, in the condensin II complex, the kleisin Cap-H2 subunit is bound by the Cap-D3 and Cap-G2 subunits (4). The functions of these eukaryotic condensin complexes are complementary: condensin I compresses chromosomes laterally and condensin II compresses them axially (6–8). Throughout much of the cell cycle, the localization of the complexes also diifers. While both complexes are critical for chromosome segregation during mitosis, in interphase, condensin I is enriched in the cytoplasm whereas condensin II predominates in the nucleus (4, 9, 10)

Condensins use ATPase activity to fuel the asymmetric extrusion of DNA loops, allowing them to disentangle chromosomes, separate homologs, and compact chromatin (11, 12). In addition to its role in cell division, condensin II contributes to the structure of interphase chromosomes. In mice and flies, condensin II antagonizes clustering of pericentric heterochromatin (7, 13, 14). Studies in *Drosophila* have also shown that the condensin II complex antagonizes the interhomolog pairing of chromosomes in somatic cells (15), and genome-wide screens have confirmed it as a central player in controlling homolog pairing behavior (13, 16, 17). The condensin II complex can thus be viewed as a master regulator of chromosome individualization, consistent with its ability to untangle and separate neighboring chromatin fibers or chromosome territories (11, 12, 18).

Intriguingly, the Cap-G2 subunit of condensin II is absent in *Drosophila melanogaster* (19). This loss in *Drosophila* is surprising, given the conservation of condensin II across most eukaryotes and its central role in essential cellular processes. *Drosophila* appear exceptional not only in their lack of Cap-G2, but also with regards to their nuclear organization. Flies align all pairs of homologs, end-to-end, in essentially all somatic tissues, a dramatic phenomenon of interchromosomal interaction not observed in any other clade (20). When pairing is observed in non-Dipteran species, it is often specific to certain genomic regions, transient, and associated with unusual or diseased states such as tumor cells in humans (21–24). In *Drosophila*, pairing of homologous chromosomes in somatic cells is critical for phenomena such as transvection, in which alleles and/or regulatory elements interact inter-chromosomally (25–27).

Given the importance of the condensin II complex in regulating pairing, we conjectured that the seemingly anomalous prevalence of pairing and the equally puzzling lack of a condensin II subunit in flies could be linked. However, it remains unclear when and how Dipterans evolved the drastic change in global nuclear organization enabling somatic homolog pairing, and it is likewise unclear if this switch to widespread pairing evolved independently or coincident with the loss of Cap-G2. Here, we investigate patterns of condensin II evolution, and the implications of these patterns for chromosome pairing in *Drosophila* and other insect species. We discovered that Dipterans are not unique in having lost a condensin II subunit. Instead, components of the condensin II complex have been repeatedly and independently lost in multiple insect lineages, with some clades missing the known condensin II-specific subunits altogether—a phenomenon not previously reported in multicellular eukaryotes. To explore the impact of the subunit losses on homologous chromosome pairing, we developed Oligopaint DNA-FISH probes (28) and quantified pairing frequencies across several insect orders with differing complements of condensin II subunits. Surprisingly, our results show that condensin II subunit losses have no relationship with pairing prevalence, and no other taxa display somatic homologous pairing to the extent seen in Dipterans. Our results rule out the intuitive idea that the loss of condensin II components and somatic homologous pairing co-evolved in Dipterans, and suggest the presence of distinct and yet undiscovered mechanisms of regulating chromosome pairing. Finally, we show that both condensin complexes and the related cohesin complex have evolved rapidly under recurrent positive selection across multiple taxa, including *Drosophila* and several mammal clades, which suggests their participation in an evolutionary arms race driven by genetic conflict. Together, our study paints a dynamic and counterintuitive view of the function and evolutionary history of condensins, and opens the door to comparative functional studies of genome organization across species.

## Results

### Multiple independent losses of condensin II subunits in insects

The Cap-G2 subunit of the condensin II complex is absent in *Drosophila melanogaster* (19), a surprising finding given that this subunit is necessary for DNA binding and is a target for the regulation of the complex in other species (29, 30). To understand when this loss occurred, we used a three-step BLAST protocol (see Materials and Methods) to search for condensin subunits, starting with Dipterans and moving outward to more distantly related insect species. We were able to identify all five condensin I subunits in virtually every species we screened, consistent with previous data suggesting that this complex should be conserved in its entirety across eukaryotes (4). When we screened for condensin II subunits, we found that all Dipterans, like *Drosophila*, are missing Cap-G2, consistent with previous reports (19). Surprisingly, Cap-G2 is also absent in several of the orders most closely related to Diptera: Lepidoptera (butterflies, moths), Trichoptera (caddisflies), Coleoptera (beetles), and Strepsiptera (twisted wing parasites) (Figure 1). The closest relatives of Diptera that retain Cap-G2 are the Hymenoptera (ants, wasps, bees) in which some but not all taxa harbor all five condensin II subunits (Figure S1). These results suggest that the Cap-G2 subunit was lost in the ancestor of Diptera over 300 million years ago, and is missing in an astonishing diversity of insects.

**Figure 1.**
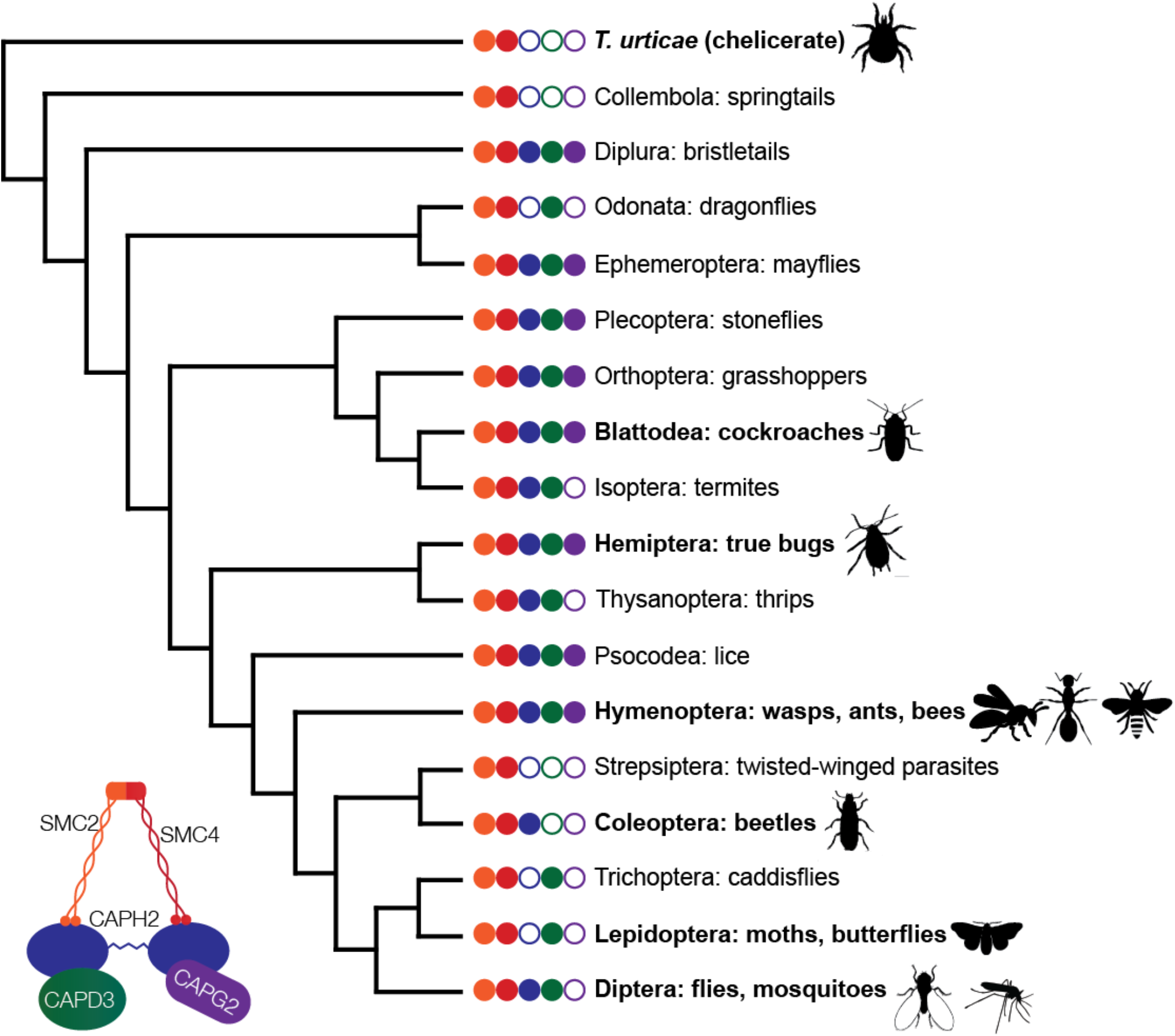
Insect phylogeny shows evidence for multiple independent losses of condensin II subunits. Based on genome sequence analysis, non-SMC subunits have been repeatedly lost in different combinations across several insect orders. Phylogeny is based on Misof et al. (62) Bold names represent orders sampled in this investigation. Condensin II subunits displayed for each order represent all subunits detected in any member of the order; some subgroups within the order have lost further subunits. In all species sampled, we were able to identify all condensin I-specific subunits. Cladogram shows species relationships only and is not to scale.

While probing these insect orders, we found that additional subunits of the condensin II complex were also missing in several taxa. For example, in addition to Cap-G2, the Cap-H2 subunit is also missing in all insects in the orders of Lepidoptera and Trichoptera, indicating a loss of this subunit approximately 200 million years ago in these lineages. In Coleoptera, Cap-D3 has been lost in all the species we screened. Even more dramatically, insects in the order of Strepsiptera are missing all non-SMC subunits of condensin II, suggesting the loss of this entire complex in some clades. Cap-H2 or Cap-D3 depletion in *Drosophila* results in lethality or sterility (15, 31), so the fact that these subunits have been jettisoned in other clades is striking.

Surprised by the losses of condensin II subunits that we observed in these orders, we expanded our study to other insect orders available in the NCBI database. Here, we found further loss events for Cap-G2, Cap-H2, and Cap-D3. Based on parsimony, many of these losses could not have originated from a single common event. Instead, each of the non-SMC condensin II subunits has been independently lost in repeated instances throughout insect evolution. We also identified more taxa where all three non-SMC subunits have been lost: Collembola (springtails), our chelicerate outgroup *T. urticae* (two-spotted spider mites), and *Trichogramma pretiosum* within Hymenoptera (Figure 1 and S1). Some of the subunit losses we identified encompassed entire orders, as in Coleoptera and Lepidoptera (Figures S2 and S3). We also found cases of more recent and dynamic losses within orders, such as the repeated losses of Cap-G2 within Hymenoptera (Figure S1), and isolated Cap-G2 and Cap-H2 loss events in Hemiptera (Figure S4). While the common ancestor of insects likely had a complete condensin II complex, the observed widespread and heterogeneous nature of condensin II subunit losses suggests that these subunits are more dispensable than previously believed. (See Figure S9 for example subunit sequences from all insect orders.)

Despite the independent nature of the condensin II subunit losses we observed, they follow a similar pattern. Among taxa missing a single subunit, Cap-G2 is missing in a vast majority of species, suggesting that this subunit is often the first to be lost, and is therefore either the most dispensable subunit or the one subject to the strongest evolutionary pressures. Species with further losses complicate existing assumptions about the roles of condensin II subunits. Coleopteran species carry Cap-H2 but are missing two subunits – Cap-G2 and Cap-D3 (Figure S2). These HEAT-repeat subunits have crucial roles in DNA binding and loop extrusion (12, 29), raising questions about the function of the condensin II complex in their absence. In Lepidoptera, Trichoptera, and Odonata, Cap-D3 is present but Cap-H2 and Cap-G2 are absent (Figure 1 and S3). Since Cap-D3 is thought to bind primarily to Cap-H2 (4, 19, 29), the nature of the association (if any) between Cap-D3 and the SMC subunits in these taxa is enigmatic.

### Condensin II loss does not induce somatic homolog pairing

The discovery of these condensin II subunit losses spread throughout insect orders offered us an opportunity to investigate the relationship between condensin II and homolog pairing. The condensin II complex antagonizes pairing in *Drosophila* (13, 15), and *Drosophila* condensin II also lacks the Cap-G2 subunit (19), raising the possibility that this loss weakens the complex’s anti-pairing function and permits heightened pairing in flies. In *Drosophila,* depletion of condensin II subunits is not well tolerated, often leading to sterility or inviability (31). Other insect species with naturally absent condensin II complexes thus offer the opportunity to study the relationship between condensin II and pairing without introducing pleiotropic effects or requiring genetic manipulation.

While it has long been believed that somatic pairing does not occur at high rates in organisms outside of Diptera (32), it has until recently been difficult to measure pairing in other species. In this investigation, we developed custom-designed, species-specific Oligopaint DNA-FISH probes (28, 33) to target unique sequences, enabling the measurement of pairing behavior at euchromatic loci in non-Dipteran insects. We chose nine insect species and a mite outgroup based on the quality of their genome assemblies, their phylogenetic positions, and the ease of obtaining live specimens. In all these species, we measured pairing levels in interphase nuclei using Oligopaints designed to 300Kb regions. (Figure 2, Figure S5 and Materials and Methods). In five species, we obtained results for two separate Oligopaint probes. Pairing levels at different loci within the same species were very similar (Figure S6), indicating that the pairing behavior we observed in each species is likely to be representative of the whole genome rather than being locus-specific. This is consistent with findings in *Drosophila* that pairing levels are similar across nearly all tissues and life stages after the early embryo (34).

**Figure 2.**
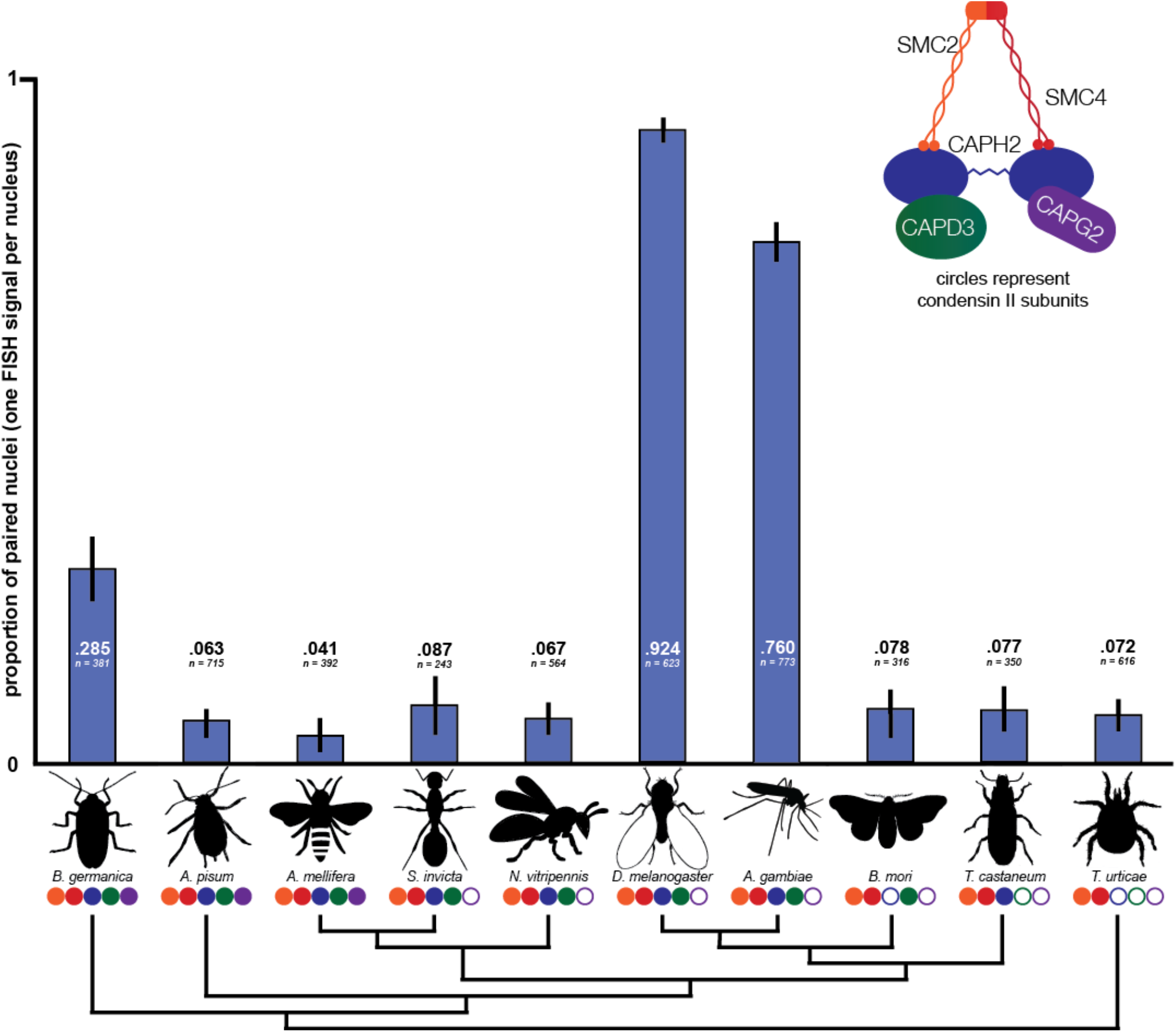
No relationship between condensin II composition and homolog pairing. Species missing more condensin II subunits do not tend to have higher pairing rates as measured by Oligopaint DNA-FISH. Blue bars represent the proportion of nuclei in each species displaying a single FISH signal. Values for each species represent the observed pairing proportion and number of nuclei scored. Each set of values represents results from a single Oligopaint probe. Error bars show 95% confidence intervals (binomial proportion with Wilson score). Circles below species names correspond to condensin II complex composition. Cladogram of insect species, based on the phylogeny of Misof et al. (62), shows relationships only and is not to scale.

In the two Dipterans that we investigated, *D. melanogaster* and *A. gambiae,* over three-quarters of nuclei displayed single FISH signals, indicating high levels of homolog pairing. However, in S. *invicta* and *N. vitripennis*, which have the same condensin II complex composition as the Dipterans, we observed low pairing levels (<10% of nuclei). This indicates that Cap-G2 loss alone does not explain the high pairing levels observed in Dipterans. Expanding our analysis to organisms with different condensin II complex composition, we found that non-Dipterans paired at lower levels regardless of the number or identity of condensin II subunits they possessed.

All non-Dipteran species displayed single FISH signals in less than 10% of nuclei, except for *B. germanica,* in which 28.5% of nuclei were paired. The outgroup chelicerate *T. urticae,* which lacks all non-SMC condensin II subunits, displayed low pairing levels, as did species like *A. pisum* and *A. mellifera* with fully intact condensin II complexes. While condensin II functions as a master regulator and potent antagonist of pairing in *Drosophila,* these results show that most non-Dipteran insect species have the capacity to keep pairing levels minimal regardless of the composition of their condensin II complexes.

### Recurrent positive selection of condensin II in flies and mammals

Since our results indicated dynamic evolution of the condensin II complex between distant taxa spanning hundreds of millions of years of evolution, we sought to understand whether these changes were accompanied by rapid evolution at the amino acid level in more closely related taxa representing shorter timescales. To understand the selective forces that drive the evolution of these complexes, we first used the McDonald-Kreitman test to detect signatures of recurrent positive selection between *Drosophila melanogaster* and its sister species *D. simulans*. In an unpolarized McDonald-Kreitman test using sequences from up to 150 *D. melanogaster* and up to 20 *D. simulans* strains, we detected strong signatures of recurrent positive selection in Cap-D2 (a component of the condensin I complex), Cap-D3 (a component of the condensin II complex), and SMC3 (a component of the cohesin complex) (Figure 3A). These results provided the first hints that the evolution of SMC complexes between *D. melanogaster* and *D. simulans* may be driven by recurrent positive selection, and are consistent with results from previous genome-wide analyses of polymorphisms in *D. simulans* (35). To detect signatures of selection in the cohesin and condensin complexes across a wider distribution of *Drosophila* species, we next used a maximum likelihood framework with Phylogenetic Analysis by Maximum Likelihood (PAML). These analyses test for recurrent changes in the sequence of a gene from a distribution of closely related species. In our sample of 17 *Drosophila* species, we found robust signatures of positive selection in nearly all components of the condensin and cohesin complexes (Figure 3B). To our surprise, almost none of the residues subject to the strongest positive selection were within putative functional domains (Figure S7).

**Figure 3.**
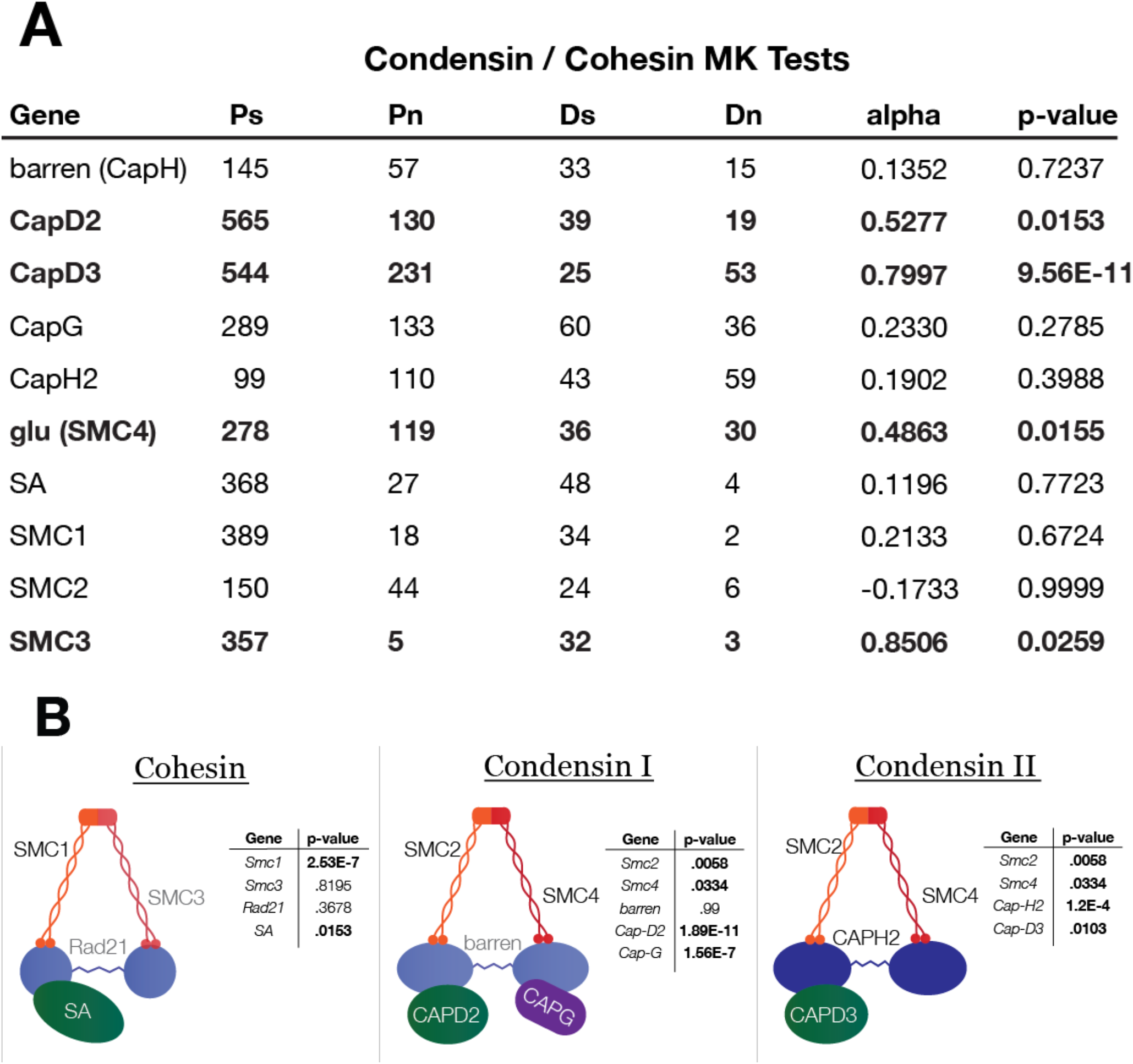
Recurrent positive selection in cohesin and condensin subunits in flies. (A) McDonald-Kreitman tests show positive selection between *D. melanogaster* and *D. simulans* in Cap-D2, Cap-D3, and SMC3. Ps = polymorphic synonymous changes within species, Pn = polymorphic non-synonymous changes within species, Ds = fixed synonymous changes between species, and Dn = fixed non-synonymous changes between species. (B) Results of Phylogenetic Analysis by Maximum Likelihood (PAML) show widespread positive selection in condensin and cohesin subunits among *Drosophila* species. P-values for PAML are derived from a log-ratio test using the log-likelihood scores for the positive selection and neutral models. Bold values represent statistically significant results.

We hypothesized that these fast-evolving residues could represent binding sites for regulators of the complex. We focused our attention on SCF^Slimb^, a ubiquitin ligase that targets Cap-H2 for degradation and has been shown to disrupt the anti-pairing activity of condensin II in Drosophila (13, 36). However, an alignment of 18 *Drosophila* species showed that the SCF^Slimb^ binding motif of Cap-H2 was well conserved, even though the amino acid sequence was often divergent across the rest of the protein (Figure S8A). This strongly suggests that the pattern of positive selection observed in Cap-H2 is not driven by pressure to escape SCF^Slimb^ regulation. Given that the knockdown of SCF^Slimb^ leads to reduced pairing (13), we also speculated that low-pairing insect species might be missing this key regulator. However, BLAST searches revealed that every species used in the pairing assay possessed a putative SCF^Slimb^ sequence, eliminating this possibility. We also found that the core binding motif for Mrg15, an important cofactor for condensin II (37), was well conserved across the same *Drosophila* species (Figure S8B). If interactions with a regulator are driving positive selection in Cap-H2, it must be a regulator with an unknown binding motif or identity.

To address whether the pattern of rapid amino acid evolution we observed in condensins is present outside of insects, we next conducted PAML analyses across 18 primate species. Our results uncovered robust signatures of recurrent positive selection in cohesins and condensins in primates as well. To ensure that the instances of positive selection we had thus far identified were not anomalous, we gathered sequences for both complexes from a variety of other mammalian clades and ran PAML on each gene in each independent clade. Our analysis shows that every one of the mammalian clades we tested has a signature of recurrent, rapid evolution for at least two genes across the complexes (Figure 4). In our analysis, we were unable to gather sufficient sequence data to analyze every subunit in some species, suggesting that the degree of rapid evolution that we observe here may be an underestimate. Together, our results suggest that despite the conserved and essential role of condensins in cell viability, the rapid evolution of these complexes is shaped by positive selection and is a general pattern across a wide variety of organisms.

**Figure 4.**
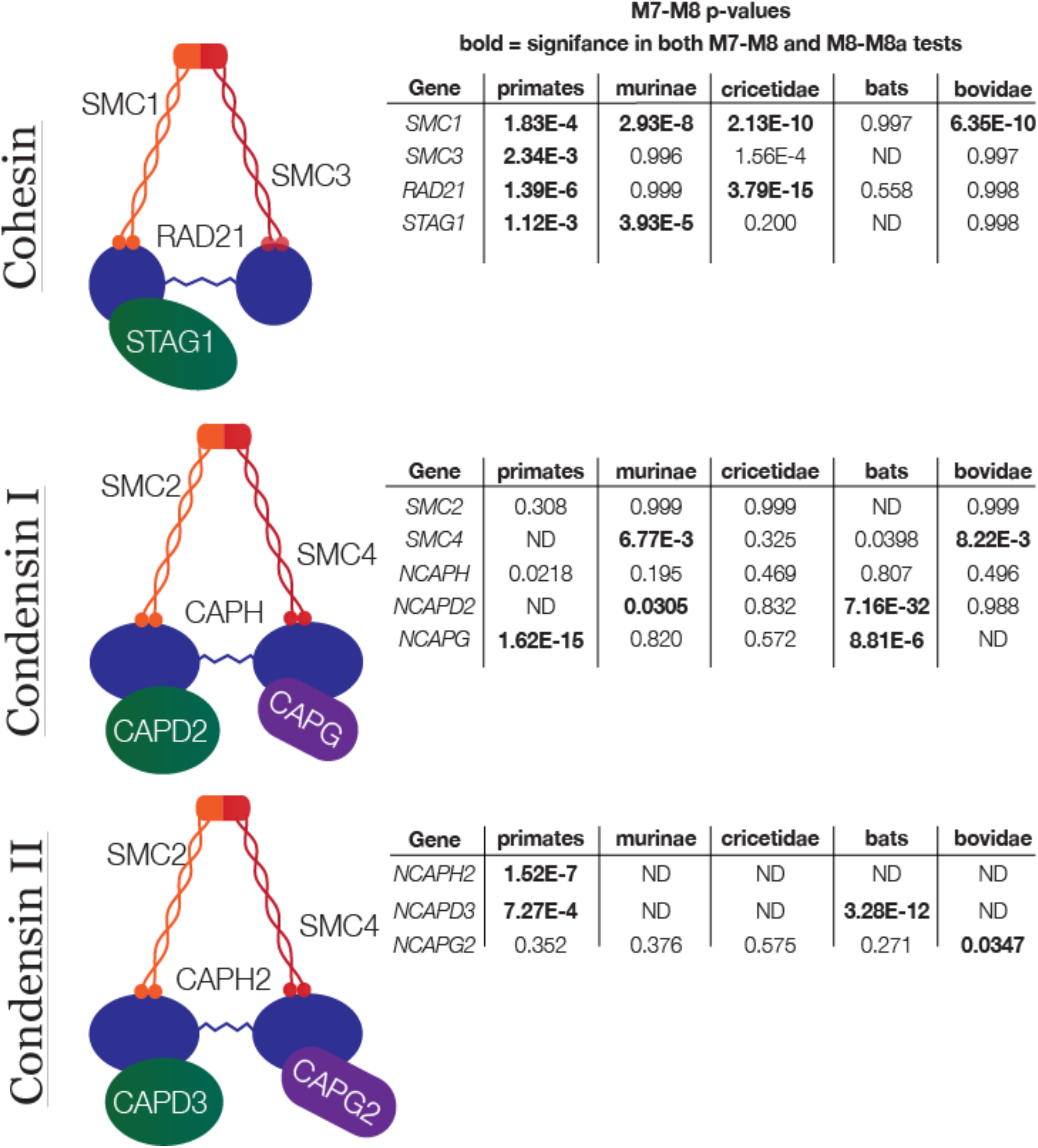
PAML analyses reveal signatures of positive selection in condensins and cohesin in mammal clades. P-values for are derived from a log-ratio test using the log-likelihood scores for the positive selection and neutral models. Bold values represent statistically significant results.

## Discussion

Condensins are ancient protein complexes that play a fundamental role in genome organization across nearly all cellular life on earth. We show that, even after billions of years of existence, condensin components evolve under recurrent positive selection, and condensin II subunits have experienced rampant losses across orders spanning hundreds of thousands of multicellular eukaryotic species. While the condensin I complex is ubiquitous in eukaryotes, the composition of the condensin II complex appears to be evolutionarily labile. The absence of the Cap-G2 subunit in Diptera initially appeared anomalous, but our results show that the Cap-G2 subunit loss dramatically predates the origin of Diptera. Nearly half of all described species on earth belong to taxa with Cap-G2 losses, indicating that the absence of a complete condensin II complex is surprisingly common. Our results further suggest that many orders of organisms are missing the condensin II complex altogether, raising questions about the mechanisms of genome organization in these taxa and across eukaryotes.

These results also raise questions regarding the consequences of condensin II component losses in these organisms. In particular, our data debunk the conjecture that condensin II complex composition directly determines the degree of pairing in an organism. Instead, we establish that somatic homologous chromosome pairing is a Dipteran-specific innovation, confirming previous speculation (32). These findings have several important implications. First, a powerful and well-established functional output of somatic pairing is gene regulation through the *trans* action of regulatory elements. To the extent that transregulation depends on close pairing of homologous chromosomes, our results predict that transvection must be limited to Dipteran species as well. Second, according to an emerging viewpoint, the pairing of homologous chromosomes may be an inevitable consequence of DNA sequence homology, and non-Dipteran species may expend considerable effort to keep homologs separate in somatic cells (38). Under this scenario, our results suggest that non-Dipteran species must utilize as yet undiscovered condensin II-independent mechanisms to separate homologous chromosomes. Alternatively, if chromosome pairing is an active process, our results raise the question of how and why such a drastic change in global nuclear organization has evolved in Dipterans.

The signatures of recurrent positive selection we observed in the condensin and cohesin complexes across *Drosophila* and mammals suggest that the evolution of these SMC complexes is shaped by evolutionary arms races driven by genetic conflict.

Previous findings regarding the role of condensins support this view. First, the condensin II complex is known to be enriched around rDNA (39) and pericentromeric heterochromatin (14). These regions are important for proper pairing and chromosome segregation in *Drosophila*, suggesting that the evolution of these complexes may be influenced by the dynamics of female meiotic drive, as is the case with centromeric histones. Second, mutants of the condensin II complex in Drosophila are viable but male sterile (4, 31), indicating their essential role in male meiosis. In *Drosophila,* chromosome decondensation is often observed in the germlines of sterile interspecies hybrid males and males carrying naturally occurring segregation distorters, raising the possibility that the condensin II complex has evolved under pressure from segregation distorters. Third, the condensin II complex has been shown to localize to retrotransposon sequences and mediate their repression in flies (40) and humans (41). As genomes continually evolve to suppress retrotransposon sequences, intragenomic conflicts involving transposable elements may also drive positive selection in Cap-D3, and potentially the rest of the condensin II complex. Finally, in addition to intra-genomic conflict, host-pathogen dynamics may also drive the evolution of condensins. In humans, SMC4 regulates the innate immune response, and in *Drosophila*, Cap-D3 responds to bacterial infection by up-regulating the expression of anti-microbial peptides (42, 43). Epstein-Barr virus, the causative agent of mononucleosis, is also known to activate Cap-G to force compaction of the host genome (44). Given the evolutionary pressures acting on the condensin complexes both from inside and outside the genome, these complexes may be positioned at the crossroads of several forms of evolutionary conflicts.

These evolutionary conflicts may not only explain the patterns of positive selection that we observed in condensin, but also the repeated loss of condensin II subunits. Though the condensin II complex plays critical roles in genome organization, other genes may have evolved to take over some of these functions when conflicts have forced the abandonment of condensin II subunits. Condensin I could simply act as a jack-of-all-trades, as it does in yeast, bacteria, and archaea. This notion is also consistent with reports of varying cellular responses to depletion of condensins across different cell types, cell lines, and species (8, 14, 45). The specific mechanics of these substitutions are enigmatic. For example, the Cap-H2 subunit of condensin II plays an integral role in chromosome territory formation in *Drosophila* (7, 18), raising the question of what, if any, alternative genome organization methods have evolved in taxa lacking this subunit. It is also possible that non-orthologous machinery may replace some condensin II functionality when it is lost. If such redundancy exists, it seems more reasonable that some lineages could jettison condensin II subunits entirely to side-step evolutionary conflicts.

Taken together, our work shows that the evolution of condensin subunits is more dynamic than previously suspected, and their function is more elastic. Our findings highlight the value of interrogating the evolution of these deeply conserved genes, and of exploring their function in non-model organisms. Despite the great heterogeneity of condensin II complex composition, we observed that non-Dipteran insects consistently organize their genomes without chromosome pairing, while Dipterans have taken a diametrically opposite path of near complete homologous chromosome pairing. Our results open the door to further investigations to elucidate the factors that underlie the mechanics of genome organization across the diversity of life.

## Materials and methods

### Inferring condensin II subunit losses

To characterize condensin II composition across *Insecta,* we sampled ~130 species, representing nearly every non-Dipteran species with a publicly available genome assembly in the NCBI database as of February 2017. For all species, we used the NCBI tBlastn function (46) to search through each insect genome, following a three-step protocol. First, we gathered publicly available sequences for all condensin I and II subunits in *D. melanogaster* (downloaded from FlyBase, based on the Release 6 genome) and *H. sapiens* sequences (from UniProt, assembly GRCh38.p12). Where multiple isoforms existed, we always chose the canonical isoform. We searched with both *D. melanogaster* and *H. sapiens* sequences against both nucleotide collection (nr/nt) and whole-genome shotgun contigs (wgs) for each target species. (For Cap-G2, we initially used *H. sapiens* sequence only.) This initial search yielded predicted condensin II subunit sequences for species within most insect orders. Within each order, we selected subunits from at least one species with a high-quality genome assembly and annotation to use as a secondary bait for species within that order. For orders where we were unable to identify annotated genes corresponding to specific subunits, we used sequences from the most closely related orders. Finally, in cases where a species had a putative subunit loss but had within-order relatives that retained the subunit, we used the subunit sequence for the most closely related species as a tertiary bait to probe the target species genome. If any of these three steps yielded a hit, we deemed the subunit to be present, and if all three failed to produce a hit, we considered the subunit to be absent. Hits yielded by this method were generally robust, with low E-values and multiple regions of homology, and we validated weaker hits by reciprocal BLAST using the sequence against annotated genomes of related species and confirming that our sequence aligned with the expected subunit. (See Figure S10 for accession numbers of identified subunits in example species.)

While demonstrating the presence of a subunit is straightforward, definitively proving the absence of one is impossible. However, several lines of evidence indicate that our three-step BLAST workflow (i.e. database sequence, within-order type sequence, and closest relative sequence if necessary) yielded robust and reproducible results. Most importantly, the condensin I complex functioned as a reliable control, because we expected its subunits not to be lost in any species. In all species we analyzed, we were readily able to identify all five condensin I subunits, suggesting that our method was able to identify subunits where they were present. Further, in many clades, we sampled multiple species and found identical condensin II complex composition in a phylogenetically consistent manner, despite substantial variation in genome assembly quality. For example, all 11 species investigated within Apoidea had lost only Cap-G2, all 16 Formicoidea species had every subunit represented, and Cap-H2 and Cap-G2 are missing in all of the 20 Lepidopteran species sampled (Figures S1 and S3). Even when condensin II complex composition within a clade was not uniform, the putative losses occurred in a phylogenetically consistent manner, thus providing further confidence in our conclusions.

### Tests for Positive Selection

Initial evidence for positive selection among condensin and cohesin subunits in *Drosophila* came from an unpolarized McDonald-Kreitman (MK) test using up to 150 *D. melanogaster* sequences and up to 20 *D. simulans* sequences obtained from PopFly (47). We found that the rate of nonsynonymous substitutions (dN/dS) between *D. melanogaster* and *D. simulans* is significantly elevated over the null expectation in these subunits: SmC3 in cohesin, Cap-D2 in condensin I, and Cap-D3 in condensin II. To validate these results, further analyses for signatures of positive selection were conducted according to the method described in Cooper and Phadnis (48). Briefly, we analyzed rates of evolution for all condensin and cohesin complex subunits in 17 species of *Drosophila,* 16 species of primates, and 6-14 species within other mammal clades. We conducted analyses for signatures of positive selection using Phylogenetic Analysis by Maximum Likelihood (PAML), and tested for recurrent positive selection by comparing NSsites models M7 (neutral) and M8 (positive selection) with 0 as the branch model. We present the p-value of the log-ratio test using the log-likelihood scores for the two models.

### Insect Husbandry

#### *Acyrthosiphon pisum* (pea aphid)

Aphids were obtained from Carolina Biological Supply Company (Burlington, NC) and maintained at 21 °C on fava bean *(Vicia faba)* plants. Slides were prepared using whole heads of adults.

#### Anopheles gambiae

Live mosquito larvae were obtained from Michael Povelones (UPENN) and maintained at 21°C in tap water. Slides were prepared from brains and head tissue of third- and fourth-instar larvae.

#### *Apis mellifera* (Western honeybee)

Bees were sourced from BeeWeaver Apiaries (Navasota, TX) and were maintained at 21 °C and provided with honey and water. Slides were prepared from dissected brains and head tissue of adult females.

#### *Blattella germanica* (German cockroach)

Cockroaches were obtained from Carolina Biological Supply Company (Burlington, NC) maintained at 28°C, and provided with water, potato slices, Blue brand dog treats, and Purina brand dry kitten food. Slides were prepared from brains and head tissue of adults.

#### *Bombyx mori* (silkworm)

Silkworms were obtained from Mulberry Farms (Fallbrook, CA), maintained at 28°C, and fed Powdered Silkworm Chow (Mulberry Farms). Slides were prepared from brains and head tissue of third- and fourth-instar larvae.

#### Drosophila melanogaster

All flies used were from a standard *w^1118^* stock, and were maintained at 21 °C on standard *Drosophila* media. Slides were prepared from a mixture of imaginal discs and brains of third-instar larvae to enrich for diploid nuclei.

#### Nasonia vitripennis

*Nasonia* cultures were grown with *Calliphoridae* pupae as a host and maintained at 18 °C. Slides were prepared from whole heads of adult females.

#### *Solenopsis invicta* (fire ant)

Ants were obtained from a wild Florida colony, maintained at 28°C, provided with water, and fed with silkworms, aphids, *Drosophila,* and Purina brand dry cat food. Slides were prepared from brains and head tissue of adult females.

#### *Tribolium castaneum* (red flour beetle)

Beetles were obtained from Carolina Biological Supply Company and maintained at 21 °C in a 20:1 mix of white flour and brewer’s yeast with carrot pieces. Slides were prepared from brains and head tissue of adults.

#### *Tetranychus urticae* (two-spotted spider mite)

Mites used were from the *Sh-Co* strain, which were isolated by Richard Clark while imbibing coffee from the Sugarhouse Coffee Company, SLC, UT (therefore named *Sh-Co).* Mites were maintained at 25 °C on kidney bean *(Phaseolus vulgaris)* and fava bean *(Vicia faba)* leaves. Slides were prepared from whole female juveniles and adults.

### Oligopaint probe design

Oligopaint probes were designed to each species using the OligoMiner pipeline (33). In brief, we retrieved genomic assemblies or contigs from NCBI Genome database, and built genome indices using Bowtie2 (Figure S10). Default settings of the OligoMiner scripts were used to mine these sequences for oligos, except for changing the length requirement to 50mers. FISH targets were chosen based on 300 kb windows with the highest density of oligos.

### Slide preparation and FISH protocol

DNA-FISH was conducted according to a protocol adapted from Larracuente and Ferree (49). Tissues specified in the preceding section were dissected in 1X PBS at room temperature. For all species, tissues were dissected from living or freshly sacrificed animals, and each batch used tissues from at least three animals. In *Drosophila,* cells from life stages and tissues other than early embryo display similar pairing levels (34), so we did not rigorously control the developmental stage or tissues analyzed between species, provided that the tissues were diploid. Immediately after dissection, tissues were placed in PBST (PBS w/ 0.1% (v/v) Triton) on ice for 5-30 minutes. Tissues were then incubated for 5-10 minutes in 0.5M sodium citrate on ice. Fixation was performed for 5 minutes directly on poly-L-lysine-coated slides using between 20 μl and 40 μl of fixative solution composed of 3:1 45% acetic acid solution and fresh 16% paraformaldehyde solution (Life Biotechnologies, Carlsbad, CA). Coverslips siliconized with SigmaCote (Sigma-Aldrich, St. Louis, MO) were pressed onto the slides firmly by hand to squash the tissues. Slides were then immediately immersed in liquid nitrogen. After immersion, coverslips were removed, and slides were incubated in cold 100% ethanol for 5 minutes. Slides were then allowed to air-dry completely. FISH hybridization buffer (FHB) was composed of 50% (v/v) 2X SSCT (SSC w/ 0.1% Tween) and 50% formamide with 20% (w/v) dextran sulfate, and stored at 4 °C. For each round of FISH, 1 μl of 10 mg/ml RNase A solution and1 μl and 3 μl of each Oligopaint probe (depending on probe reliability) were added to 25-40 μl of FHB (depending on tissue volume). The complete FHB and probe mix was then placed on coverslips and added to the slides. Slides were then incubated on a heat block at 90.7 °C for 7 minutes, sealed with Parafilm, and placed overnight in a humidified chamber at 37 °C. After hybridization, slides were washed in 2X SSCT for 15 minutes, in 0. 1X SSC for 15 minutes, and in 0.1X SSC with 0.1% (v/v) DNA stain (either DAPI or TOPRO-3) for 15 minutes. Slides were then air-dried, mounted with Vecta-Shield media (Vector Laboratories, Burlingame, CA), and sealed with nail polish.

### Microscopy and quantitative measures of homolog pairing

All images were acquired using a Zeiss LSM 880 Airy Scan confocal microscope. Airy scan function was not employed. For all species, at least 5 images were acquired from areas enriched for diploid cells displaying strong signal. In *B. mori, S. invicta,* and *T. castaneum*, two separate slides were imaged and scored. Images were processed using ImageJ software and scored manually. Nuclei were scored as having zero, one, two, or three or more FISH signals, and a separate category for grossly polyploid nuclei was tracked. Multiple signals were judged to exist if multiple peaks of FISH intensity, in tandem with multiple clearly discernible shapes, were distinguishable by eye, even if the edges of these signals overlapped. The pairing proportion was defined as the number of nuclei displaying one signal divided by the sum of one-signal and two-signal nuclei. Slides or regions containing high proportions of zero-signal or abnormal nuclei were excluded from analysis.

## Acknowledgements

This work was supported by the National Institutes of Health R01 GM115914 (NP), R35 GM128903 (EJ), (Genetics Training Grant 5T32GM007464-40 (CJL), a Mario Capecchi endowed chair in Biology (NP), and the Pew Biomedical Scholars Program (NP). We are grateful to those who provided us with biological materials: Dale Clayton and Scott Villa for pigeon lice, Richard Clark for spider mites, Rebecca St. Pierre and Jack Longino for fire ants, Nels Elde for bees, Jack Werren for Nasonia wasps, Michael Povelones for Anopheles mosquitoes, and T. DeLonge for creatures from above.

---

**Figure S1.**
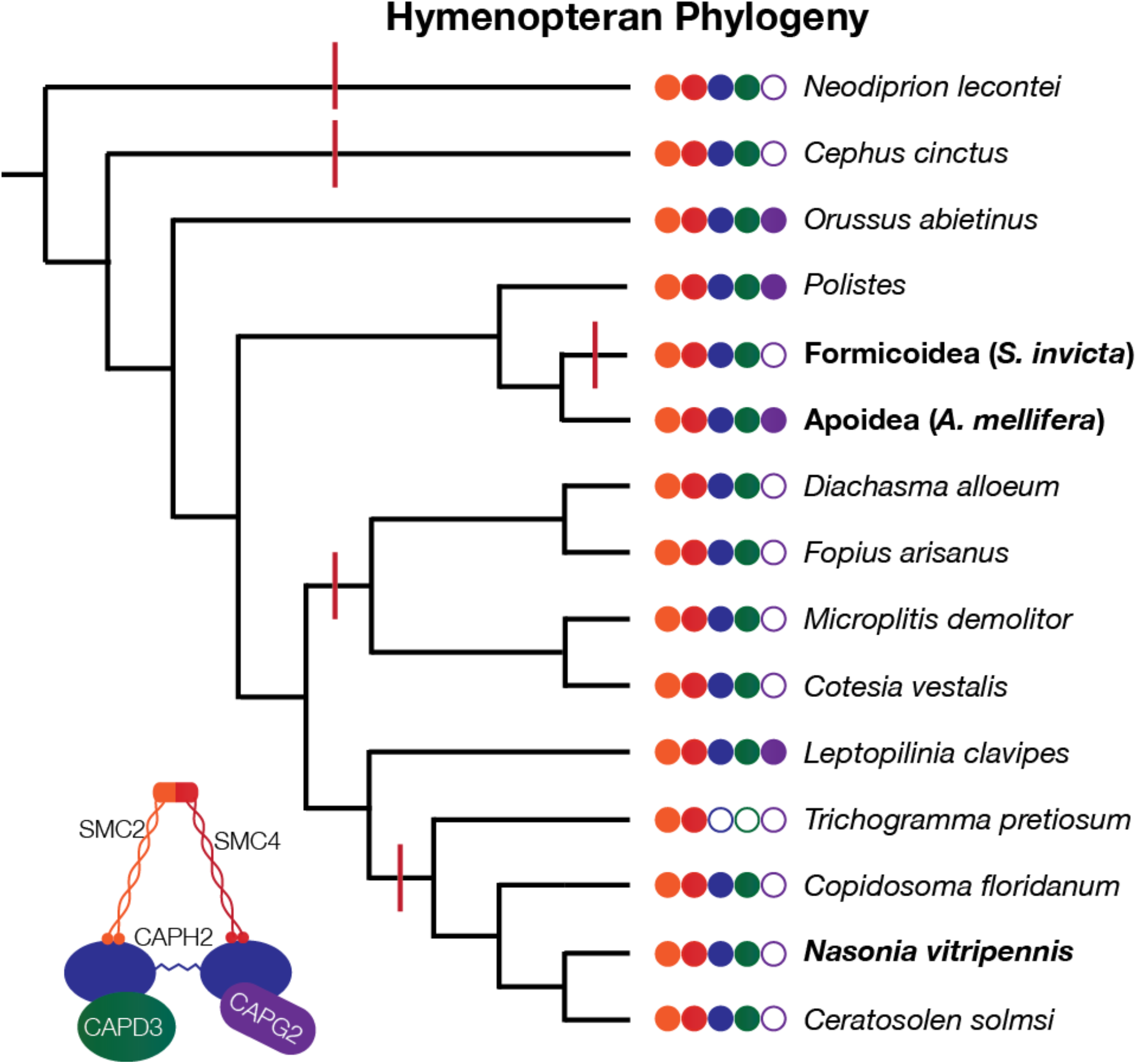
Hymenopteran phylogeny shows independent within-order losses of CapG2. Species phylogeny is based on Peters et al. (50) with distinctions within Ichneumonoidea based on Quicke and van Achterberg (51). Bold names represent clades sampled in this investigation. Red lines indicate inferred independent losses of condensin II subunits. All condensin I subunits are present in all species shown. Cladogram shows species relationships only and is not to scale.

**Figure S2.**
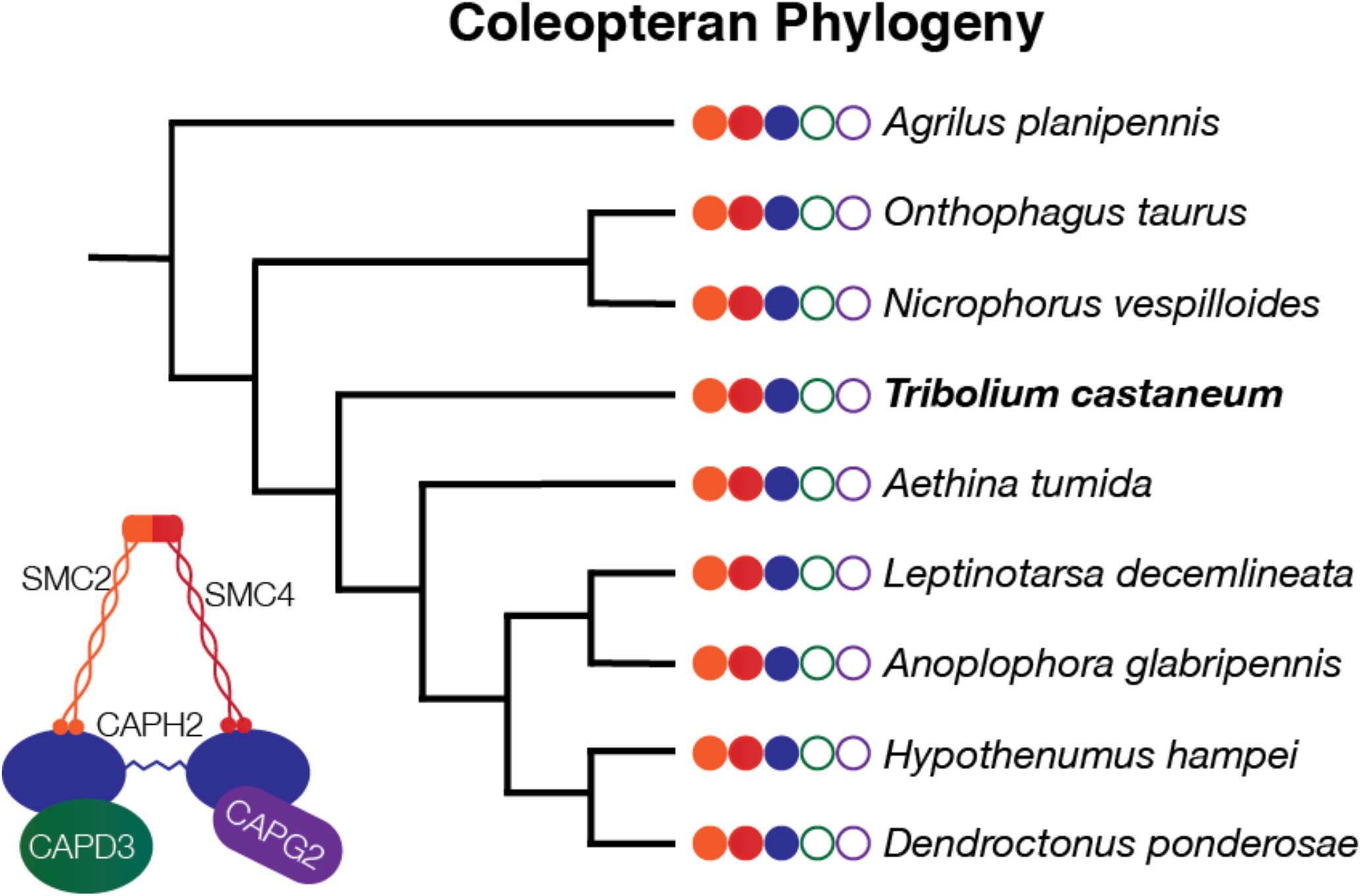
Coleopterans show consistent patterns of CapD3 and CapG2 loss. All condensin I subunits are present in all species shown. Cladogram is based on the phylogeny of Zhang et al. (52), shows species relationships only, and is not to scale.

**Figure S3.**
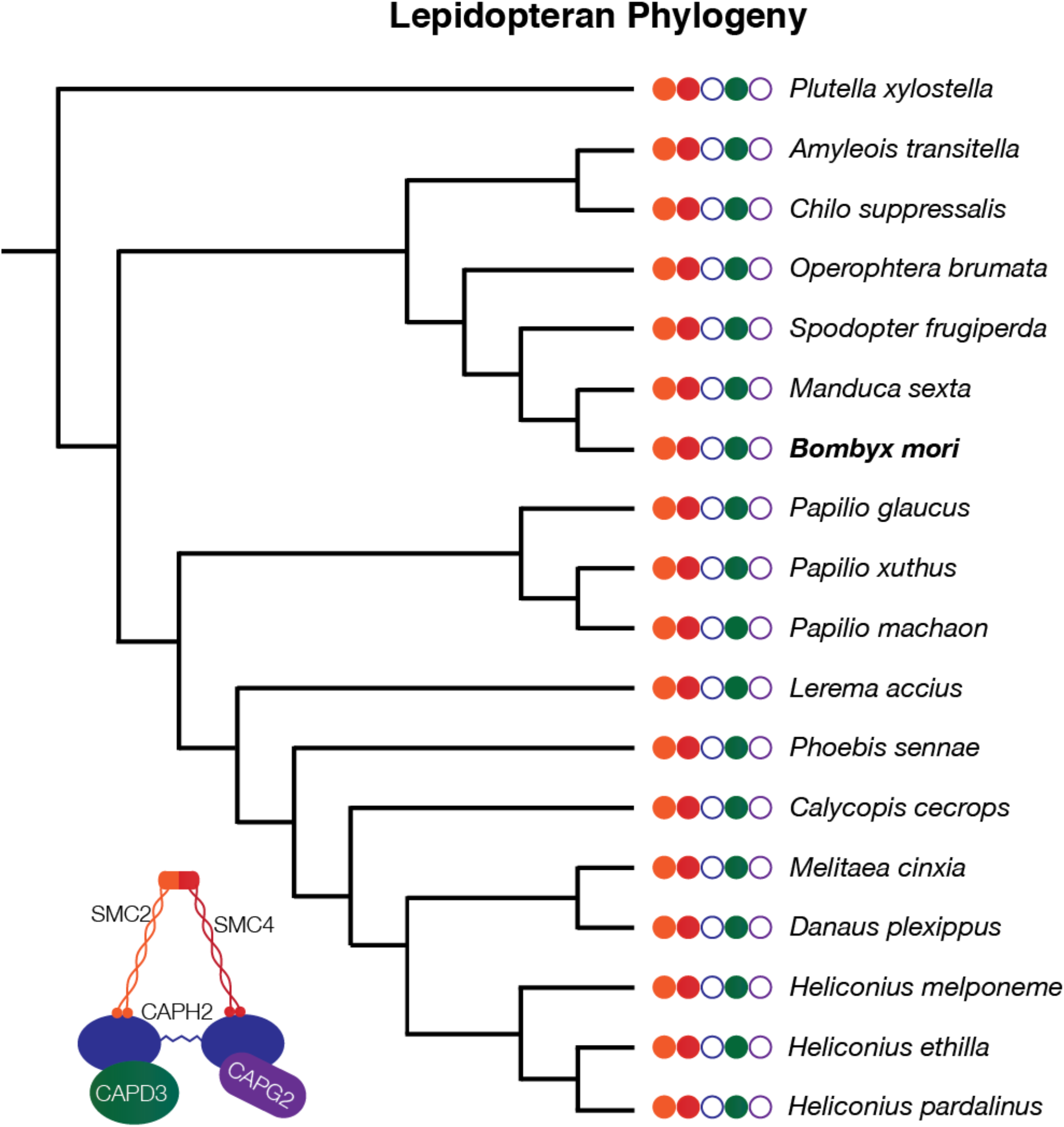
Lepidopterans show consistent CapH2 and CapG2 loss. Whole-order phylogeny based on Regier et al (53), with Nymphalidae based on Freitas and Brown (54), Heliconius based on Kozak et al. (55), and Papilio based on Zakharov et al. (56). All condensin I subunits are present in all species shown. Cladogram shows species relationships only and is not to scale.

**Figure S4.**
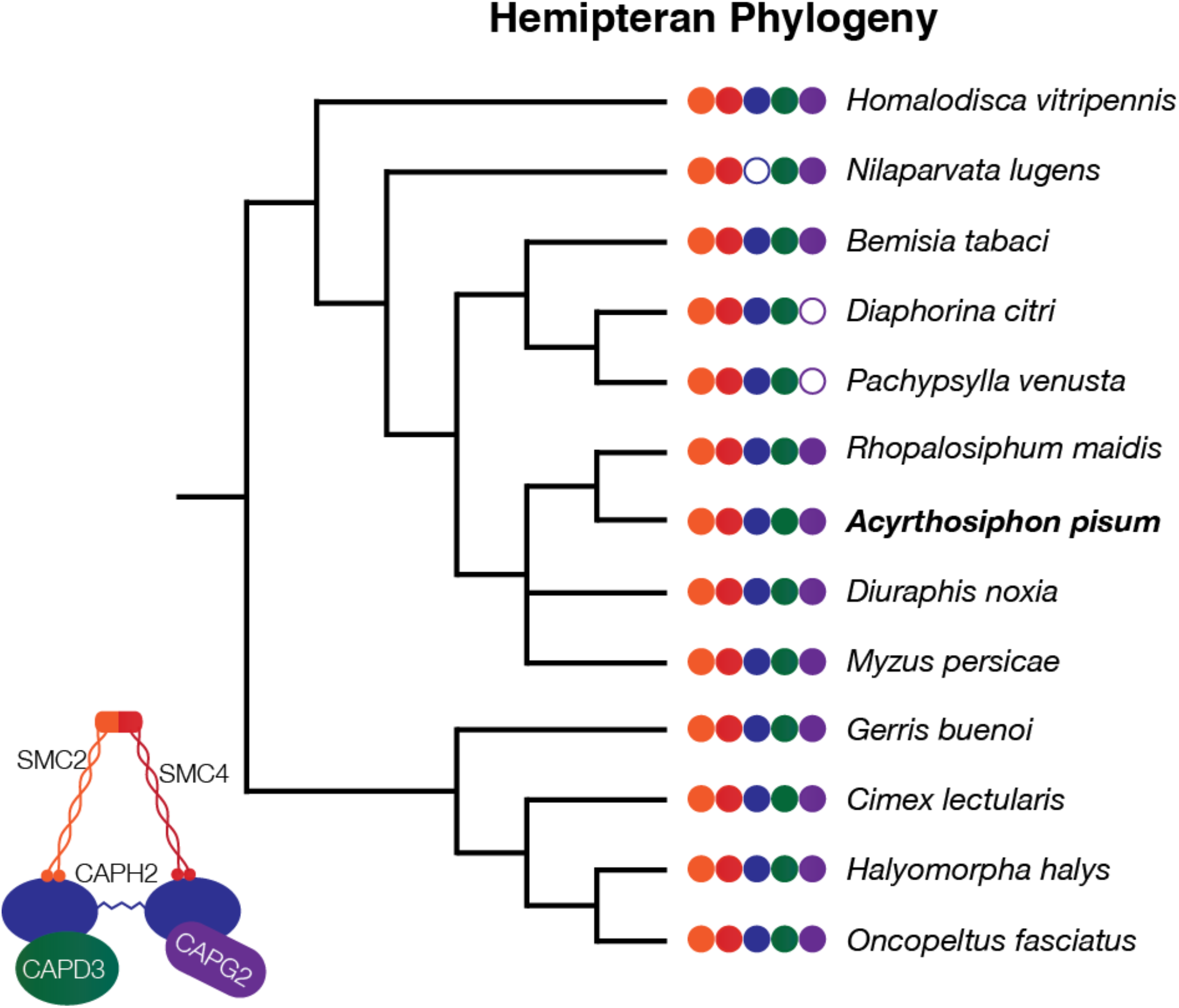
Hemipteran phylogeny shows independent loss events of CapG2 and CapH2. All condensin I subunits are present in all species shown. Cladogram is based on the species phylogeny of Song et al. (57), with distinctions within Aphididae based on Nováková *et al.* (58), and is not to scale.

**Figure S5.**
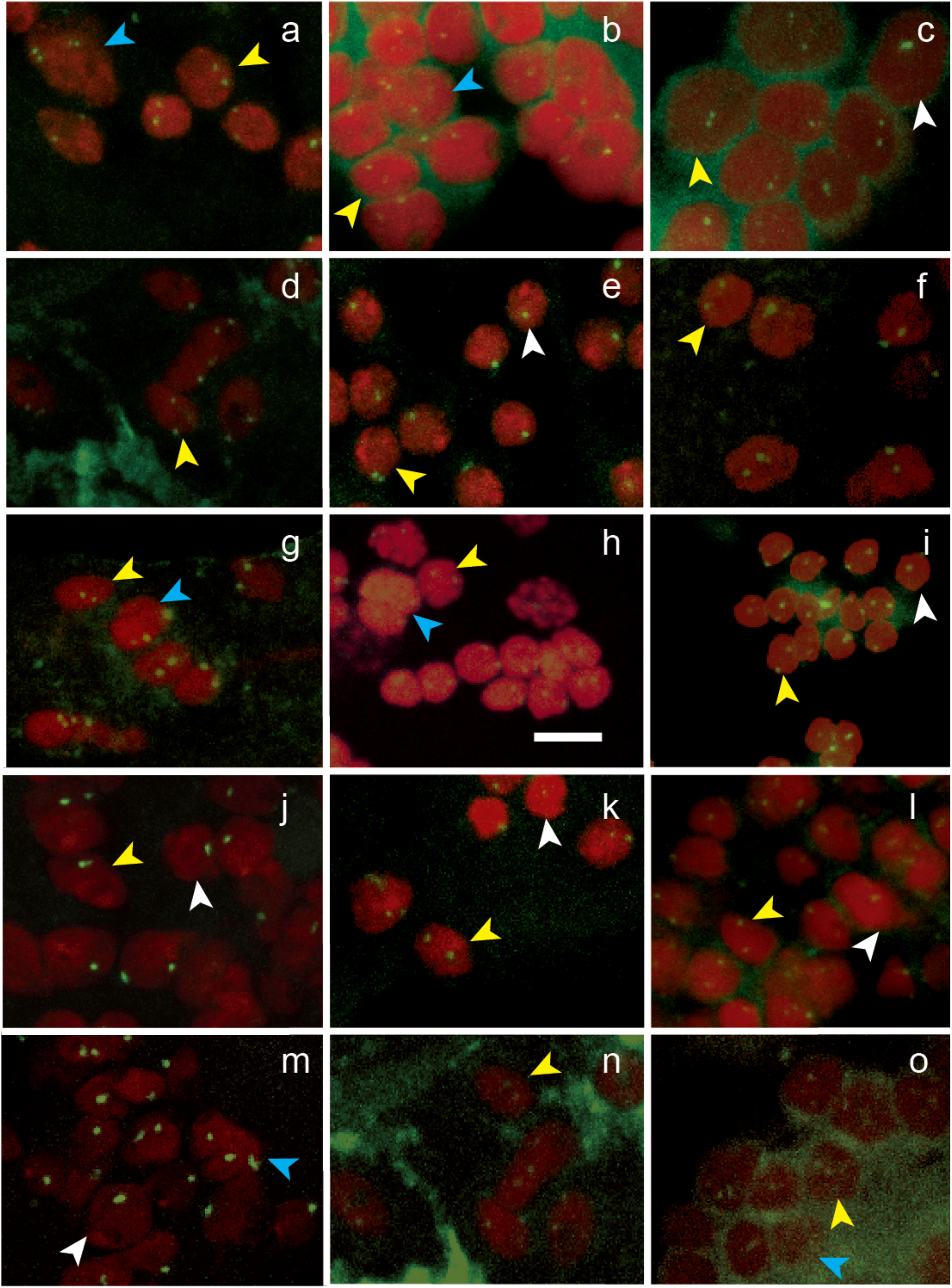
Representative FISH images. For all images, red represents DNA stain, and green represents the Oligopaint probe channel. All images are at the same scale, and the scale bar in panel H represents 5 μm. Images A-J are from the main probes used for each species, and images K-O show results from the second probe, when one was used. DNA in images A-E and I-N is stained with DAPI, and DNA in images F-H and O is stained with TOPRO-3. Image A shows the probe NP17, labeled with Cy3, in *A. mellifera* tissue. Image B shows NP01 (Cy3) in *A. pisum,* C shows NP04 (Cy5) in *B. germanica,* D shows NP05 (Cy3) in *B. mori,* E shows Null4 (Cy5) in *D. melanogaster,* F shows NP09 (Cy3) in *N. vitripennis,* G shows NP11 (Cy3) in S. *invicta,* H shows NP13 (Cy3) in *T. castaneum,* I shows NP16 (Cy5) in *T. urticae,* and J shows 231 (Cy3) in *A. gambiae.* For alternate probes, K shows NP18 (6-FAM) in *A. mellifera,* L shows NP02 (Cy5) in *A. pisum,* M shows 232 (Cy5) in *A. gambiae,* N shows NP06 (Cy5) in *B. mori,* and O shows NP10 (6-FAM) in *N. vitripennis.* In all images, white arrows indicate examples of nuclei scored as paired (if present), yellow arrows indicate examples of unpaired nuclei, and blue arrows indicate examples of nuclei excluded from calculations of pairing proportions due to apparent polyploidy, lack of signal, indeterminate nuclear boundaries, or poor signal-to-noise ratio (see Materials and Methods).

**Figure S6.**
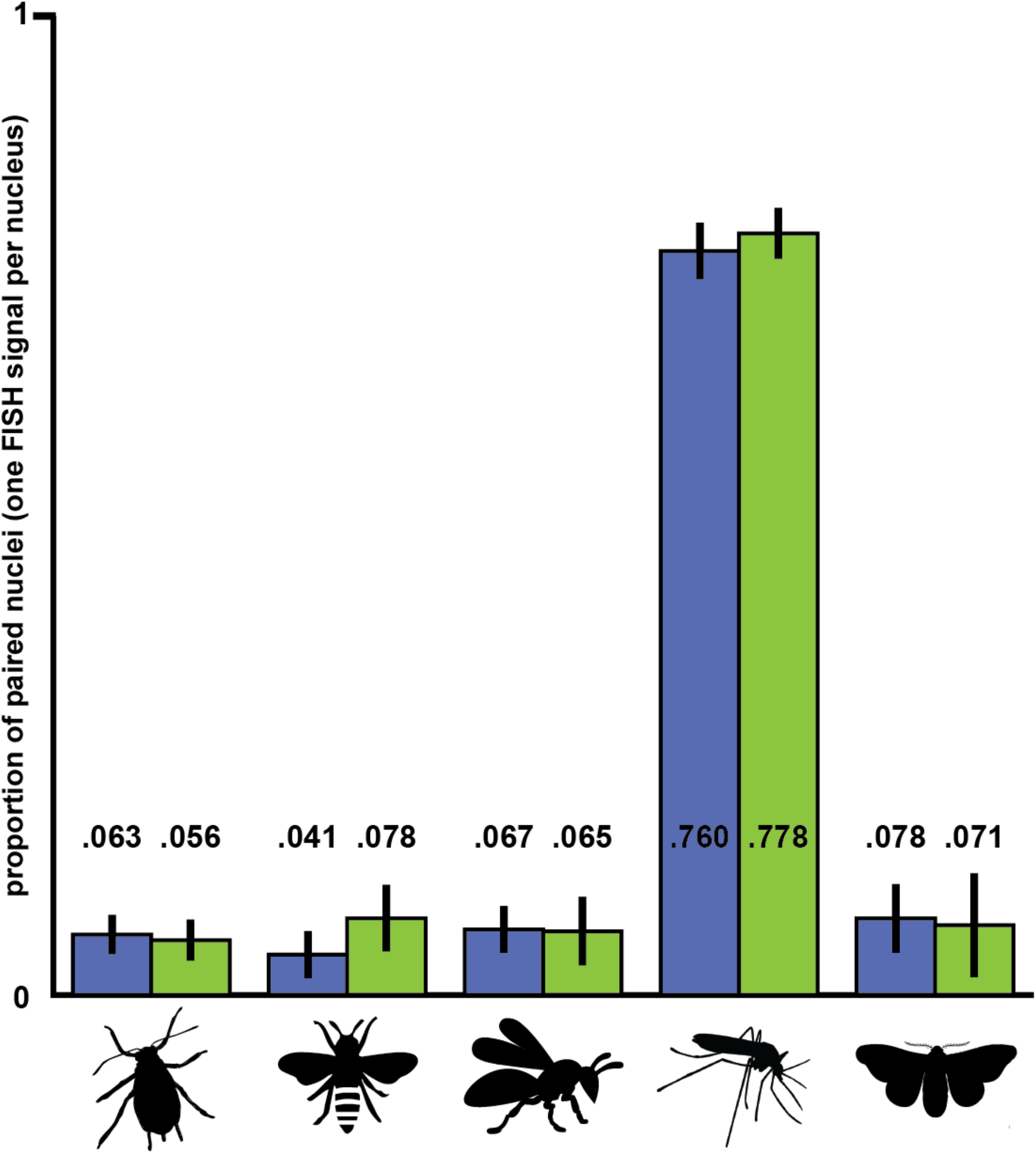
Pairing data for alternate probes indicates little within-species variation. In *A. pisum, A. mellifera, N. vitripennis, A. gambiae,* and *B. mori,* results were obtained for both Oligopaint probes synthesized. Results for the probes with worse signal or fewer nuclei scored in each species were excluded from the main analysis, but are reported here. Main probe results are shown in blue, and alternate probe results in green. Proportion of nuclei with single FISH signals observed with the alternate probes was highly consistent with that observed in main probe, except in *A. mellifera.*

**Figure S7.**
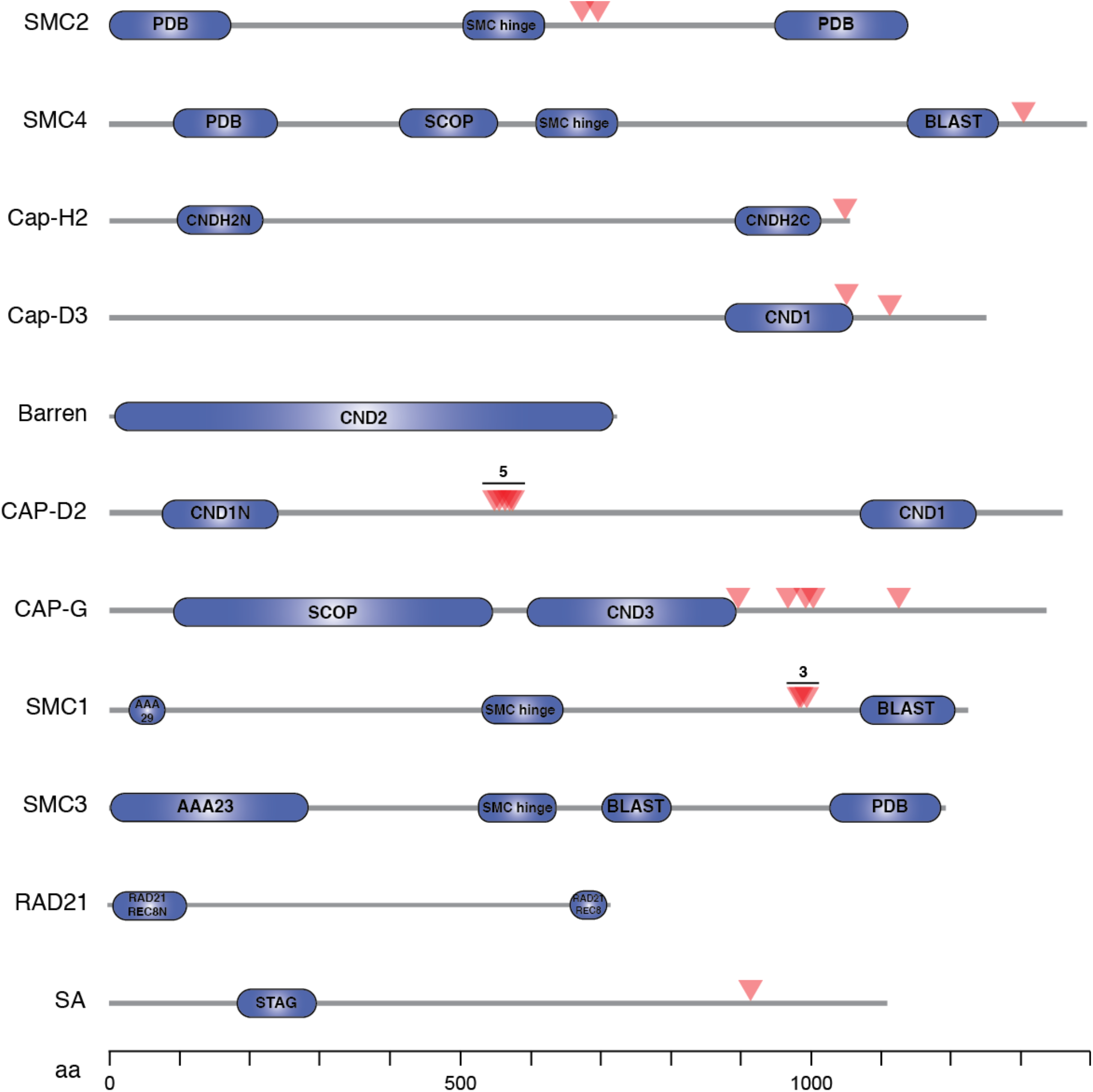
Sites of repeated positive selection in condensin and cohesin subunits. Red arrows show residues subject to positive selection (dN/dS) across insect lineages, as indicated by PAML. Boxes indicate putative functional domains as identified by SMART (59).

**Figure S8.**
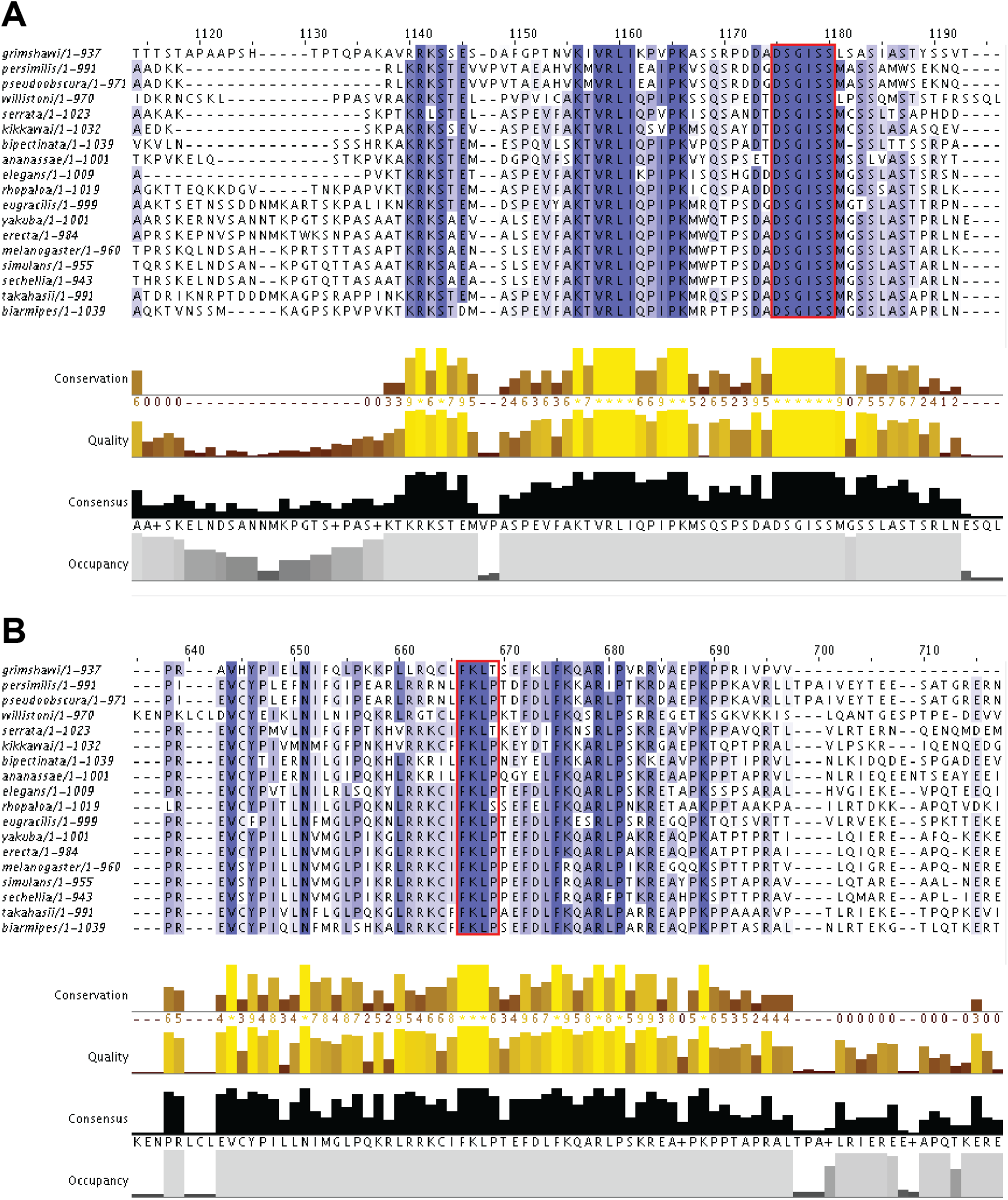
Cap-H2 regulator binding motifs show strong conservation across *Drosophila.* Cap-H2 sequences of 18 *Drosophila* species representing ~40 million years of evolutionary divergence were aligned with MUSCLE software and visualized with Jalview (60, 61). Darker blue represents more highly conserved residues. (A) We found high conservation relative to the surrounding region in the SLMB binding motif (DSGISS, outlined in red). (B) We also observed high conservation in the Mrg15 binding motif (FLKP, in red).

